# Somatosensory cortex and body representation: Updating the motor system during a visuo-proprioceptive cue conflict

**DOI:** 10.1101/2024.09.23.614575

**Authors:** Jasmine L. Mirdamadi, Reshma Babu, Manasi Wali, Courtney R. Seigel, Anna Hsiao, Trevor Lee-Miller, Hannah J. Block

## Abstract

The brain’s representation of hand position is critical for voluntary movement. Representation is multisensory, combining visual and proprioceptive cues. When these cues conflict, the brain recalibrates its unimodal estimates, shifting them closer together to compensate. Converging evidence from research in perception, behavior, and neurophysiology suggest that such updates to body representation are communicated to the motor system to keep hand movements accurate. We hypothesized that primary somatosensory cortex (SI) is crucial in this updating process due to its role in proprioception and connections with primary motor cortex. We tested this hypothesis in two experiments. We predicted that proprioceptive, but not visual, recalibration would be associated with change in short latency afferent inhibition (SAI), a measure of sensorimotor integration (influence of sensory input on motor output) (Expt. 1). We further predicted that modulating SI activity with repetitive transcranial magnetic stimulation (TMS) should affect recalibration of the proprioceptive estimate of hand position, but have no effect on the visual estimate or on the normal inverse relationship between proprioceptive and visual recalibration (Expt. 2). Our results are consistent with these predictions, supporting the idea that (1) SI is indeed a key region in facilitating motor system updates based on changes in body representation, and (2) this function is mediated by unisensory (proprioceptive) processing, separate from multisensory visuo-proprioceptive computations. Other aspects of the body representation (visual and multisensory) may be conveyed to the motor system via separate pathways, e.g. from posterior parietal regions to motor cortex.

**Significance Statement:** Representation of the hand, which is critical for accurate control of movement, comes from weighting and combining available proprioceptive (position sense) and visual cues. Our results suggest that when the hand representation is modified by introducing a conflict between these cues, the motor system receives updates directly from the primary somatosensory cortex (SI). These updates are specific to the change in proprioceptive representation and are absent when cues are not in conflict. This may represent a unisensory pathway to the motor system that conveys information about hand representation, acting in parallel with multisensory pathways involving posterior parietal and premotor regions.

## INTRODUCTION

Performing complex movements with the hands is a universal human experience. By adulthood we have learned countless such motor skills: tying our shoes, controlling a computer mouse, etc. Understanding how the brain learns in these situations is complicated by the involvement of multiple motor, sensory, and multisensory processes. “Learning” likely entails not only improving the speed-accuracy tradeoff (motor skill) and adjusting motor commands to compensate for any systematic movement errors (motor adaptation), but also fine-tuning perception of the hand itself. The brain may adjust how it interprets proprioceptive (position sense) cues from the muscles and skin, recalibrating its proprioceptive estimate of hand position in response to sensory prediction errors^1^. Importantly, if we can see our hands, then our brain’s estimate of hand position is likely multisensory, arising from a weighted average of both visual and proprioceptive cues^2,3^. If visual and proprioceptive cues conflict but are still interpreted as arising from the same source (the hand), the brain is expected to recalibrate each unimodal estimate of hand position in proportion to its relative variance (Fig. 1A), updating the integrated multisensory estimate of hand position^2^.

**Fig. 1.**
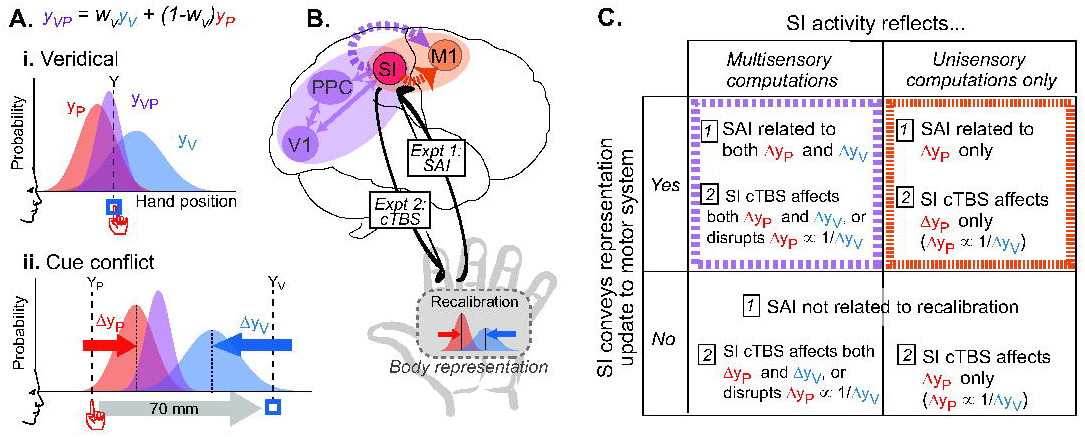
Models and predictions. **A.** Multisensory hand representation (y_VP_) is generally conceived as the result of multisensory integration, a weighted sum of the unimodal visual (y_V_) and proprioceptive (y_P_) estimates, with weight of vision vs. proprioception (w_V_) determined by various environmental and internal factors^2,29,30^. **i.** Even when visual and proprioceptive cues of hand position (blue square and red hand) match, i.e. are veridical, unimodal estimates (y_P_ and y_V_) may disagree slightly, due to differences in the processing pathways of each sensory modality^31,32^. Multisensory integration provides a single estimate of hand position to guide movement (y_VP_). **ii.** When a spatial conflict arises between visual and proprioceptive cues, the brain is thought to unconsciously recalibrate (shift) each unimodal estimate (Δy_P_, Δy_V_) closer together, reducing the apparent mismatch^2^. The magnitudes of Δy_P_ and Δy_V_ are inversely related, with the lower-weighted modality usually recalibrating more^2,9,14^. **B.** Visuo-proprioceptive recalibration likely arises from interactions among multisensory regions of posterior parietal cortex (PPC) and unisensory visual and somatosensory regions (e.g., SI). Such updates in hand representation must be transmitted to frontal motor regions (M1) to keep movement accurate. We hypothesize this process to involve SI, either conveying multisensory information (purple dashed arrow) or proprioceptive information alone (orange arrow). Expt. 1 assesses the effect of a change in hand representation (recalibration) on the SI-M1 pathway, measured with short latency afferent inhibition (SAI). Expt. 2 modulates activity in SI, M1, or sham with continuous theta burst transcranial magnetic stimulation (cTBS), assessing the impact on hand representation. **C. Predictions.** Expt. 1 could show that in the presence of cue conflict, SAI is either related to both Δy_P_ and Δy_V_ (purple cell), or to Δy_P_ but not Δy_V_ (orange cell), which would indicate SI activity reflects either multisensory computations (first column) or proprioceptive computations only (second column), respectively. Either result would suggest that SI conveys hand representation updates to the motor system (top row). Alternatively, SAI might not be related to recalibration, which would indicate SI does not convey such updates to the motor system (bottom row). Expt. 2 could show that SI cTBS affects both Δy_P_ and Δy_V_, or the inverse relationship between them (first column, i.e. SI activity reflects multisensory processing) or that it affects Δy_P_ but not Δy_V_ (second column, i.e. SI activity reflects only the proprioceptive component of recalibration). Alternatively, SI cTBS could affect neither modality of recalibration, which might suggest SI is involved only in lower level aspects of proprioceptive processing. The results of Expt. 1 and Expt. 2 support the upper left (orange) cell, i.e. SI conveys hand representation updates to the motor system in terms of proprioceptive recalibration specifically.

Visuo-proprioceptive recalibration in hand perception is an example of the brain’s ability to update body representation. Converging lines of evidence from research in perception^4,5^, behavior^6–8^, and neurophysiology^9,10^ suggest that such updates to body representation are communicated to the motor system to keep hand movements accurate. For example, even in the absence of motor learning, recalibration resulting from visuo-proprioceptive cue conflict altered hand movements^11^. Perceptual recalibration has also been associated with changes in primary motor cortex (M1) excitability. We used transcranial magnetic stimulation (TMS) to test M1 excitability before and after participants experienced conflicting or veridical visuo-proprioceptive information about index finger position^9^. No performance feedback was available, and participants recalibrated both proprioceptive and visual estimates of the index finger as expected^12–15^. Results indicated that M1 excitability decreased in association with proprioceptive recalibration, but increased in association with visual recalibration^9^. These M1 changes were somatotopically focal, affecting only the index finger representation and not forearm or biceps representations^10^.

The observed modality-specific changes in M1 excitability could suggest that the neural mechanism conveying body representation updates to the motor system involves plasticity in areas traditionally considered unisensory, such as primary somatosensory cortex (SI)^16,17^ or early visual areas that have indirect connectivity with M1^18^. SI, encompassing areas 1, 2, 3a, and 3b, is an intriguing candidate region. SI is linked both with (1) parietal regions likely involved in multisensory visuo-proprioceptive perception^19–24^, and (2) frontal motor regions, including M1 (Fig. 1B). The degree to which somatosensory input affects motor output, termed sensorimotor integration, can be measured neurophysiologically: Short latency afferent inhibition (SAI) describes the inhibitory effect of a peripheral electrical stimulus delivered before a motor response evoked by transcranial magnetic stimulation (TMS)^25^. The exact neuronal pathway is not known, but thought to be mediated at the cortical level by projections from SI pyramidal cells to M1 interneurons^26,27^.

Here we asked two questions about the function of SI (Fig. 1C): (1) Is SI important for conveying hand representation updates to the motor system, and (2) If so, is the involvement of SI multisensory, involving both visual and proprioceptive updates, or unisensory (related to proprioception only)? In Experiment 1 we assessed SAI before and after 22 participants estimated the position of a series of veridical or conflicting visual and proprioceptive cues about their left index finger (Fig. 3A). Each participant experienced both conditions in random order. In Experiment 2, three groups of 27 participants received continuous theta burst TMS (cTBS) over the SI or M1 representation of their left index finger, or sham (Fig. 4A). cTBS is thought to reduce cortical activity for up to an hour^28^. All participants estimated the position of a series of veridical visuo-proprioceptive cues about their left index finger before and after cTBS, followed by the introduction of a visuo-proprioceptive cue conflict.

## METHODS

### Participants

A total of 121 self-reported right-handed healthy young adults participated in the study. All participants had normal or corrected-to-normal vision, and no contraindications to TMS ^33^. Participants self-reported that they were free of neurological, musculoskeletal, attention, or learning conditions. Procedures were approved by the Indiana University Institutional Review Board. All participants provided written informed consent before participating in the study.

### Perceptual alignment task general methods

Expt. 1 and 2 used variations of the same behavioral task to examine participants’ perceived hand position using visual, proprioceptive, or bimodal cues. In addition, participants in both experiments were asked to rate their level of attention, fatigue, and pain from TMS at the conclusion of each session to evaluate whether different sessions or groups had similar subjective experiences. Participants were also questioned about their perceptions of the bimodal target to evaluate whether they perceived an offset during cue conflict. Details of the session and trial block design, which differ between experiments, are explained in their respective sections.

<u>Apparatus (</u>Fig. 2A). Participants were seated in front of a custom 2-D virtual reality apparatus with a two-sided infrared touchscreen (PQ Labs) and a mirror positioned at eye level. The touchscreen was positioned beneath the mirror such that visual information appeared in the plane of the touchscreen. A fabric drape attached to the apparatus was fastened around their neck, preventing vision of their limbs or surrounding environment. The participants always kept their right hand above the touchscreen and their left hand below the touchscreen.

**Fig. 2.**
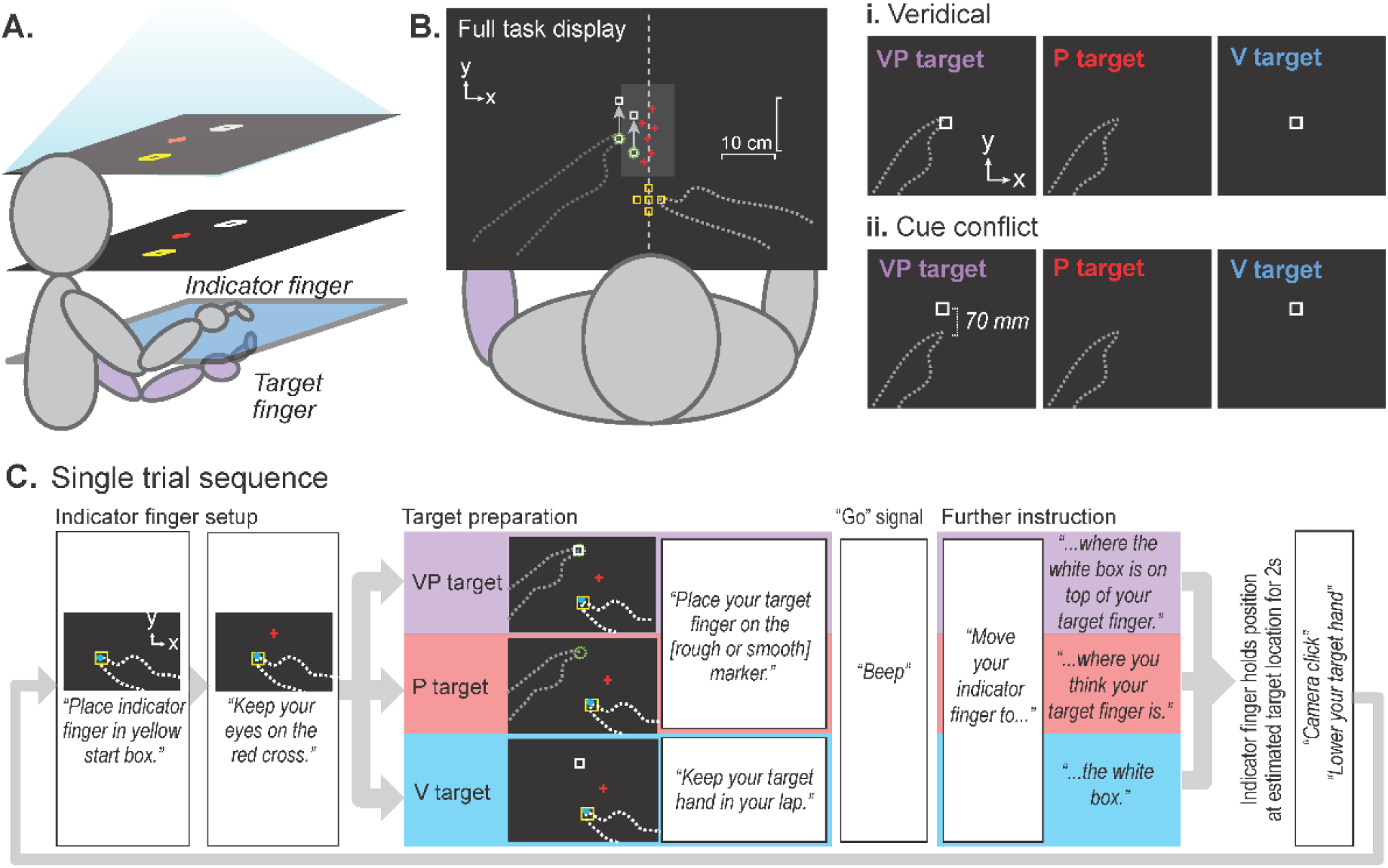
Visuo-proprioceptive alignment task. **A. Apparatus.** Participants viewed task display in a horizontal mirror (middle layer), making images appear to be in the plane of the touchscreen (bottom layer). Participants used their right index finger (indicator finger) on top of the touchscreen to indicate perceived target positions related to the left (target) index finger below the touchscreen. Participants had no direct vision of their hands at any time and upper arms were covered with a drape. **B.** Indicator finger starting positions varied randomly among 5 locations centered at the body midline (yellow squares). Targets varied between two positions slightly to the left of body midline (green circles), about 30 cm in front of the body and well within reach. Participants were asked to gaze at a randomly-positioned red cross within a zone along the midline, rather than at any visual targets (white squares). **i-ii.** The three target types, presented in pseudorandom order, were indicated by both visual and proprioceptive cues (VP target), proprioceptive cues only (P target), or visual cues only (V target). Target fingertip on a tactile marker beneath the touchscreen provided proprioceptive cues, and a white square provided visual cues. Not to scale. Dashed lines not visible to subjects. No performance feedback was ever available. During cue conflict blocks (ii), the white square gradually shifted forward imperceptibly for both VP and V targets, 1.67 mm per trial, to a max of 70 mm. **C.** Single trial sequence. Each trial began with audio instructions for positioning the indicator finger, on top of the touchscreen, in the yellow start box. A blue cursor appeared during this phase, only until the indicator finger was correctly positioned. Instructions for positioning the target finger beneath the touchscreen came next, followed by the “go” signal. If the indicator finger did not leave the start box after 3s, further audio instructions were played to remind the subject what to do. When the indicator finger had left the start position, touched down in a new position (without dragging), and held still for 2s, the endpoint position was recorded and a camera click sound played to let the subject know the trial was over.

<u>Target types (</u>Fig. 2B). Participants were asked to use their unseen right index fingertip (indicator finger) to indicate the position of one of three target types: a visual only target (V), a proprioceptive only target (P), and a visuo-proprioceptive target (VP). For the V target, participants were asked to indicate their perceived position of a projected visual white square (1 cm). For the P target, participants were asked to indicate their perceived position of their left index fingertip, which was placed on one of two tactile markers below the touchscreen. For the VP target, participants were asked to indicate their perceived position of the white box projected on top of their left index fingertip. All subjects were told explicitly that the white box would always be right on top of their target fingertip on VP target trials. For V target trials, participants rested their left hand in their lap.

<u>Single trial procedure (</u>Fig. 2C). Each step of the trial was cued with recorded audio instructions. The trial began with the participant placing their indicator fingertip in a yellow square that served as the starting position. The starting position was presented at one of five locations centered at the body midline, about 20 cm in front of the chest. A blue cursor representing their indicator fingertip was initially shown to help position their fingertip in the starting square, which then disappeared to avoid any feedback cues. Next, participants were asked to fixate a red cross that appeared at random coordinates within a 10 cm zone along the body midline. They were asked to try to keep their gaze there throughout the remainder of the trial to reduce the chances that gaze behavior might differ between different target types. Participants were then instructed where to place their target hand (in lap for V targets, on one of the tactile markers for P and VP targets). The tactile marker (1 cm) used in the P and VP targets was one of two positions (3 cm apart), resulting in 10 start position-target position pairs, requiring different directions and extents of movement by the indicator hand. The purpose of this was to make it less likely that participants would simply repeat the same motion with their indicator hand rather than trying to estimate the target position.

The proprioceptive cue of P and VP targets was achieved by having participants actively position their own left finger on the tactile marker. They had to lower their hand to their lap between each trial, and it remained in the lap for V trials. The purpose was to make sure that every P and VP trial involved the subject recently moving their own hand, which is thought to reduce proprioceptive drift^34^. Recent active movement may also improve proprioceptive salience, so that across participants reliance on visual vs. proprioceptive cues will be approximately even. In the present study and previous studies, this procedure results in people relying equally or slightly more on proprioception^32,35^, enabling us to see the full range of possible recalibration behaviors (some people will recalibrate vision more than proprioception and vice versa).

When both hands were correctly positioned, participants heard a go signal and, at their own pace, lifted their indicator finger off the touchscreen and placed it down where they perceived the target to be. If they hesitated longer than 3 seconds, further instructions played to remind them what to do (Fig. 2C). There was no speed requirement or knowledge of performance or results. Adjustments were permitted, and the trial terminated after the indicator fingertip remained still on the touchscreen for 2 seconds. The x-y coordinates of the indicator finger endpoint served as a proxy for where the participant perceived the target to be.

### Experiment 1 design

#### Participants

Twenty-two adults (12 male, 21.6 ± 3.68 years, mean ± SD) participated in Experiment 1. Participants completed 2 sessions each. In the Cue Conflict session, a 70 mm mismatch between visual and proprioceptive cues about the left index fingertip was gradually imposed in the sagittal plane. In the Veridical session, visual and proprioceptive information remained veridical. The sessions were separated by at least 5 days, with order randomized, to minimize carry-over effects. On average there were 16 ± 14 days (mean ± SD) between sessions. TMS measurements were performed pre- and post-perceptual alignment task in each session (Fig. 3A).

**Figure 3.**
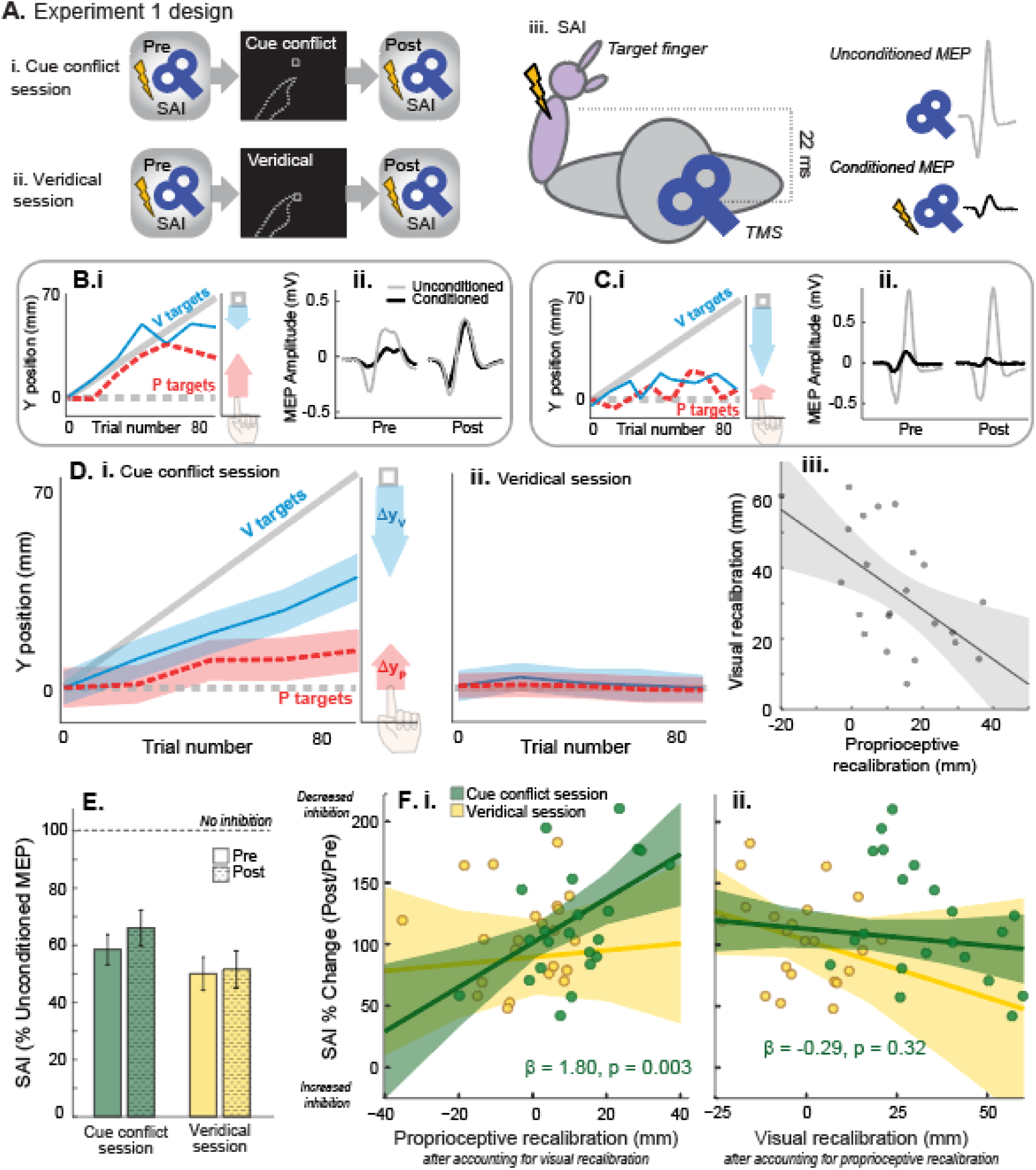
Expt. 1 design and results. **A.** Experiment 1 design. All participants completed two sessions in random order. SAI was assessed immediately before and after the behavioral portion of each session. **i.** Cue conflict task consisted of V, P, and VP trials, with the visual cue gradually shifting forward relative to the proprioceptive cue. **ii.** The Veridical session included the same number of V, P, and VP trials, with cues displayed veridically throughout. **iii.** SAI was assessed by delivering an electrical stimulus to the left median nerve 22ms prior to a suprathreshold TMS pulse over right M1. Top: Schematic unconditioned motor evoked potential (MEP). Bottom: Schematic of MEP conditioned by median nerve stimulus, illustrating inhibition of motor output by somatosensory input. **B-C.** Expt. 1 example participants’ performance on P and V targets in the cue conflict block (averaged every 4 trials for clarity) (i) and their corresponding changes in SAI in the cue conflict session (ii). Undershooting the visual target on V trials represents visual recalibration (blue arrow) and overshooting the proprioceptive target on P trials represents proprioceptive recalibration (red arrow) (i). **B.** Participant with relatively high proprioceptive recalibration (i) demonstrated a decrease in SAI (larger ratio of conditioned relative to unconditioned MEP) post -task relative to pre (ii). **C.** Participant with relatively larger visual recalibration demonstrated an increase in SAI (smaller ratio of conditioned relative to unconditioned MEP) post-task compared to pre. **D.** Group means on unimodal P and V trials (red and blue) during the cue conflict session (**i**) and veridical session (**ii**), averaged every 4 trials for clarity. Shaded regions represent SEM. **iii**. In the cue conflict session, individuals who recalibrated proprioception to a greater extent recalibrated vision to a lesser extent (r = -0.52, p = 0.013). **E.** Group mean SAI pre- and post-alignment task for the cue conflict and veridical sessions. SAI was expressed as the conditioned MEP amplitude relative to the unconditioned MEP amplitude (test stimulus alone). Values < 100 indicate inhibition, with smaller values denoting greater inhibition. Error bars are standard error of the mean. SAI did not differ significantly between sessions or timepoints. **F.i-ii**: Change in SAI (post divided by pre) plotted against predictor residuals, with lines of best fit and corresponding 95% CIs. In the cue conflict session, recalibration of either modality in the positive direction is beneficial (helps compensate for the visuo-proprioceptive mismatch). **i:** After statistically controlling for the effect of visual recalibration, large proprioceptive recalibration in the cue conflict session (green) was significantly associated with reduced SAI (disinhibition). There were no associations between proprioceptive recalibration in the veridical session (yellow) and SAI. **ii:** After controlling for the effect of proprioceptive recalibration, there were no significant associations between visual recalibration and change in SAI for either session.

### Behavior

#### Perceptual alignment task blocks

There were two blocks of trials in each session. The first block was identical between sessions and consisted of a baseline veridical block with 15 V, 15 P, and 10 VP trials that were pseudorandomized. The second block comprised 84 trials (42 VP, 21 V, and 21 P, alternating order). In the Cue Conflict session (Fig. 3Ai), the white box was gradually shifted forward from the target index fingertip on VP trials in the second block. The offset increased by 1.67 mm per VP trial. By the end of the block, the visual cue was displaced 70 mm forward of the proprioceptive cue, in the sagittal plane. Participants rarely become aware of this manipulation ^36^. In the Veridical session, there was no offset and VP targets remained veridical throughout (Fig. 3Aii).

#### Visual and proprioceptive recalibration

Figure 3Bi and Ci shows indicator finger estimates relative to the true positions of V and P targets for two exemplar participants across trials in the cue conflict block. Similar to past work, the magnitude of visual and proprioceptive recalibration (Δy_P_ and Δy_V_) was calculated using indicator fingertip endpoints of the V and P targets, respectively. The average y-estimate of the last first four trials of the given modality was subtracted from the average y-estimate of the first four:

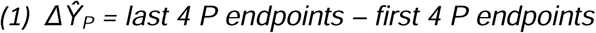

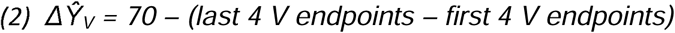

Recalibration was expressed relative to the change in true y-position of the target (70 mm for V targets, 0 mm for P targets at the end of the cue conflict block; 0 mm for both modalities in a veridical block). In the cue conflict condition, a positive recalibration value indicates recalibration in the expected direction (i.e., undershooting for V targets and overshooting for P targets).

#### Weighting

Weighting was calculated using the 2D estimated indicator fingertip position in VP targets relative to the unimodal targets during the first block of each session (veridical targets). For instance, a person that relies more on vision would be expected to have their mean estimate of VP targets closer to the mean estimate of V targets compared to P targets. Based on formula (3) a weighting value greater than 0.5 indicates greater reliance on vision whereas a value less than 0.5 indicates greater reliance on proprioception.

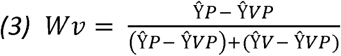

Consistent with previous work ^37^, we computed a separate Wv for each VP trial, comparing the VP trial endpoint with the 4 closest V and 4 closest P trials in the sequence. If the V and P trial endpoints were too closely overlapping (less than 0.5 SD apart), no Wv was computed for that VP trial.

#### Visual and proprioceptive endpoint variance

Variance was computed for the 2D indicator finger endpoints on unimodal V and P trials during the first block of each session (veridical). This was accomplished by converting each endpoint to a vector relative to the mean [x,y] coordinates of the cluster of endpoints. Variance was then computed for the vector magnitudes. To approximate the variance of participants’ visual and proprioceptive estimates (as opposed to the variance of the endpoints, which includes motor and proprioceptive variance from the indicator hand), we subtracted ½ P variance from each estimate^37^. This assumes that the variance contributed by the indicator hand to variance of any target type is half of the variance apparent in the indicator finger endpoints on unimodal P targets, which of course is an approximation.

### Neurophysiology

#### M1 stimulation

TMS was applied over the M1 hand representation in the right hemisphere, to probe neurophysiology pertaining to the target hand that experiences visuo-proprioceptive cue conflict. Single monophasic TMS pulses were delivered using a Magstim 200^2^ stimulator (Magstim Company LTD, UK) with a 70 mm figure-of-eight coil. The coil was held tangentially to the scalp with the handle 45 degrees postero-lateral from the midline to elicit posterior-to-anterior current in the right M1. The hotspot was identified by the scalp position that elicited the largest and most consistent response in the left first dorsal interosseous (FDI) muscle. The location and trajectory were registered in BrainSight neuronavigation system (Rogue Research, Montreal, Canada) for consistent coil positioning throughout the session. Surface electromyography (EMG) was recorded from the left FDI muscle and abductor pollicis brevis (APB) muscle using a belly-tendon montage and a ground electrode over the ulnar styloid process. EMG recordings were amplified (AMT-8; Bortec Biomedical), band-pass filtered (10-1000 Hz), sampled at 5000 Hz, and recorded using Signal software (Cambridge Electronic Design Ltd, UK). At the beginning of each session, we found resting motor threshold, defined as the minimum intensity that elicits a twitch >= 50 µV in at least 10 out of 20 trials^33^. We then found the stimulus intensity that elicited a twitch of 1 mV on average over 10 trials (SI_1mV).

#### SAI procedure

SAI was assessed before and after the perceptual alignment task in each session (Fig. 3A). To elicit SAI, a TMS pulse at SI_1mV was delivered 22 msec after an electrical stimulus at the left median nerve. Electrical stimuli were delivered with a Grass Instruments S88 stimulator (Astro-Med, West Warwick, RI) with an in-series stimulus isolation unit (SIU-5) and a constant-current unit (CCU-1) (square wave pulse, 0.2-ms duration, cathode proximal). The intensity was set based on the lowest intensity that elicited a slight thumb twitch and consistent APB M-wave amplitude. The M-wave amplitude was monitored online and kept constant throughout the session ^38,39^.

20 conditioned pulses (median nerve stimulus + TMS) and 20 unconditioned pulses (TMS alone) were delivered in a random order, with an interstimulus interval of ∼ 5 sec, both pre- and post-alignment task. We adjusted the TMS intensity post-alignment task, if needed, to elicit the same size unconditioned MEP response of ∼1 mV. Therefore, any changes in SAI reflect changes in somatosensory projections to M1 rather than changes in M1 excitability alone.

SAI magnitude was expressed as a percentage of the average conditioned MEP peak-to-peak amplitude relative to the unconditioned peak-to-peak amplitude (Fig. 3Aiii). Therefore, lower numbers of SAI within a time point (pre or post) indicate greater inhibition by the somatosensory afferent volley. We also calculated the change in SAI (post/pre) for each session. Therefore, smaller delta SAI indicates greater inhibition post relative to pre, while larger delta SAI values indicate less SAI post relative to pre.

### Statistical Analysis

Baseline (pre-alignment task) neurophysiology was compared between the veridical and cue conflict sessions using paired t-tests for RMT, SI_1mV, and SAI. To test whether the alignment task influenced SAI differently across sessions, we performed a Session (cue conflict vs. veridical) x Time (pre vs. post) repeated measures ANOVA.

Multi-level hierarchical modeling was used to assess whether an individual’s magnitude of proprioceptive or visual recalibration was related to their change in SAI. We first examined a full model that included predictor variables for the interaction between Session and Modality, and their main effects. However, the full model had variance inflation factor (VIF) values that suggested the presence of multicollinearity (VIF value of 6). Therefore, we computed a reduced model that only had the interaction term of Session and Modality, which had VIF values that ranged between 1.04-1.05. Since the reduced model had less multi-collinearity and was not statistically different from the full model (Chi-squared test: p = 0.7), the reduced model was used for analysis. Finally, we computed Pearson’s correlation coefficients between baseline SAI (pre-alignment task) and 2D Wv in the veridical session, and between baseline SAI and P variance and V variance in the veridical session. We also computed the correlation between baseline SAI and proprioceptive recalibration in the cue conflict session. Alpha was set at 0.05.

### Experiment 2 design

#### Participants

81 individuals (38 male, 21.68 ± 4.32 years, mean ± SD) participated in Experiment 2. All were right-handed according to the Edinburgh Handedness Inventory^40^. Participants were assigned to one of three stimulation groups: SI, M1, or Sham (N=27 per group). Group assignment was random, following a block randomization table. Group assignment was revealed to the experimenters only when it was time to deliver cTBS, after task training and baseline measures were complete. SI had 12 male participants (21.44 ± 3.94 years, mean ± SD), M1 had 13 male participants (21.3 ± 4.58 years, mean ± SD), and the Sham group had 13 male participants (22.3 ± 4.28 years, mean ± SD). Sample size was determined *a priori* with a power analysis on pilot data in earlier participants who received SI, M1, or no cTBS (N=8,13,27) with the same session design. We determined a total sample size of 81 would be needed to have 80% power to detect an effect size f of 0.404 with α of 0.05 with a 1-way ANOVA on 3 groups. The effect size was determined for proprioceptive recalibration in the pilot subjects.

Continuous theta burst stimulation (cTBS) was delivered in between two veridical perceptual alignment blocks (Fig. 4A) to assess any effect of cTBS on baseline behavior. This was followed by a cue conflict block. The second veridical block and the cue conflict block took about 45 minutes for most subjects to complete, such that all subjects finished the behavioral tasks within an hour of receiving cTBS.

**Figure 4.**
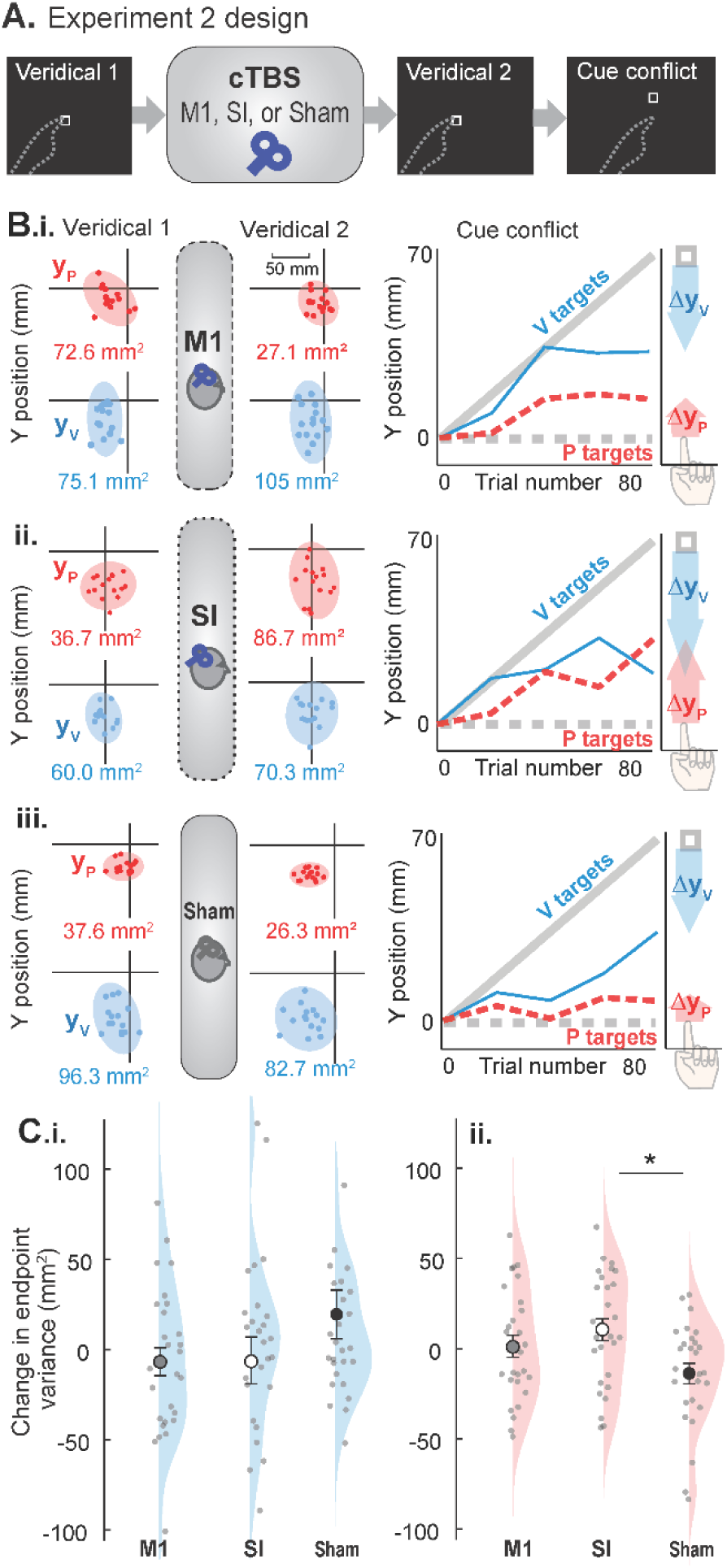
**A.** Experiment 2 design. After a baseline block of V, P, and veridical VP trials, participants received M1, SI, or sham cTBS according to random group assignment. After a second block of veridical trials, participants performed a cue conflict block in which the visual cue gradually shifted forward relative to the proprioceptive cue. **B. i-iii.** Example participants’ indicator finger endpoints in 2D on P (red) and V (blue) unimodal trials before and after cTBS (veridical 1 and 2) for M1, SI, and sham groups, respectively. Data plotted with target always at the origin. Participant was seated in the direction of the negative y-axis. Shaded ellipses represent 95% confidence intervals, whereas red and blue numbers represent 2D variance calculated as described in the Methods. Each person’s performance during the cue conflict block is also displayed, with shaded arrows representing recalibration in visual (blue) and proprioceptive (red) estimates. **C.** Group mean changes and standard errors on unimodal V (i) and P trials (ii) from Veridical 1 to Veridical 2. Dots represent individual participants. **i.** There was no effect of group on change in visual variance. Not pictured: one outlier in the Sham group at 332 mm^2^, and one outlier in the SI group at -243 mm^2^. **ii.** SI cTBS increased proprioceptive variance relative to sham cTBS. All participants are visible in this axis range.

### Behavior

The apparatus, target types, and task instructions were identical to that of Experiment 1. Each participant performed three blocks of perceptual alignment task trials: Veridical 1, veridical 2, and a cue conflict block, with cTBS delivered after Veridical 1 (Fig. 4A). Each veridical block consisted of 40 trials in repeating order (15 V, 15 P, 10 VP). After the second veridical block, participants completed the same cue conflict block used in Experiment 1. Recalibration and weighting were computed as in Experiment 1. Endpoint variance was computed as in Experiment 1, except we did not subtract 0.5 P target variance from each variance estimate because the planned analyses were a pre-post design.

In order to test whether cTBS over SI had any effect on tactile sensitivity that could interfere with participants’ ability to feel the tactile markers with their target finger, we performed an abbreviated grating orientation test (GOT) before and after cTBS for all participants. I.e., the first GOT was performed after Veridical 1 and before cTBS, and the second GOT was performed after cTBS and before Veridical 2. We used J.V.P. DOMES for cutaneous spatial resolution measurements. The set consists of domes of 35, 0.5, 0.75, 1.0, 1.2, 1.5, 2, 3, 4, 5 mm grating width. For the assessment, participants had their eyes closed and their left hand resting on the table with the palm facing up. They were tested on their left index fingertip palmar surface. The domes were attached to a force gauge to maintain uniformity in the pressure (0.65N – 0.95N) applied while testing based on an earlier study ^41^. The testing started with a 3mm grating width and a random order of orientation (Vertical or Horizontal) was used. No feedback was provided to the participant. If the subject correctly reported six orientations consecutively then a grating of smaller width was used next. If they reported one or more orientations incorrectly then a larger grating was used. This continued until they were unable to report the orientation correctly. The smallest grating width that participants reported correctly was noted as their tactile sensitivity threshold.

### Transcranial Magnetic Stimulation

TMS was delivered using a using a Magstim Super Rapid Plus stimulator with a D70^2^ 70-mm figure-of-eight coil (Magstim Company LTD, United Kingdom). Single pulses were first delivered over right M1 to find the resting motor threshold of the left FDI muscle using identical methods as Experiment 1. Brainsight Neuronavigation was used as in Experiment 1, with participants’ heads registered to the template brain for consistent coil positioning throughout the session.

cTBS was delivered over the right M1 or right SI to modulate the hemisphere that pertains to the left hand (target hand), which experiences the visuo-proprioceptive cue conflict. For the M1 group, cTBS was delivered over the FDI motor hotspot. The SI stimulation target was defined 2 cm lateral and 1 cm posterior to the M1 target^42–44^. Given the limitations on depth of stimulation in a flat coil^45^, regions within SI most likely stimulated are areas 1, part of 2, and part of 3b; area 3a is likely too deep in the central sulcus to be stimulated effectively with this method^46^. The sham target was the same as the M1 target, but consisted of an unplugged coil placed on the scalp. A second coil, held behind the participant’s head out of their view, was plugged in so that the sounds of the cTBS train would be audible.

cTBS was delivered at 70% of RMT ^44,47,48^, consisting of triplets of 50 Hz repeated at 5 Hz for 40 seconds ^28^. Participants sat quietly for five minutes before and after cTBS to avoid any potential reversal of the after-effects ^49^.

### Statistical Analysis

To determine whether cTBS affected endpoint variance on unimodal P and V targets, subtracted individual values for Veridical 2 minus Veridical 1 and ran a 1-way ANOVA on the difference across groups. We analyzed weighting of vision vs. proprioception the same way (Wv). Two participants had closely overlapping distributions of P and V endpoints in one of the veridical blocks and insufficient Wv were calculated to form an estimate for the whole block.

These participants were excluded from the Wv analysis. ANOVAs were followed by Tukey’s HSD post-hoc comparisons upon significant result. Each distribution was checked for normality and equality of variances (Shapiro-Wilk test and Levene’s test, respectively). Only the change in V target variance violated any assumptions (due to an outlier in the Sham and SI groups), so the non-parametric equivalent (Kruskal-Wallis) was used instead.

We tested for group effects on P, V, and total recalibration (sum of P and V recalibration, to approximate the total magnitude participants compensated for the 70 mm conflict) using separate 1-way ANOVAs, which were followed by Tukey’s HSD post-hoc comparisons upon significant result. GOT thresholds pre-cTBS were subtracted from post-cTBS and these changes were compared across groups using the Kruskal-Wallis test. Αlpha was defined as 0.05 for all hypothesis tests.

### Control experiment design

#### Participants

18 individuals (4 male, 22.22 ± 4.41 years, mean ± SD) participated in the control experiment. All were right-handed according to the Edinburgh Handedness Inventory^40^.

### Behavior

Because the main experiments used the same method to introduce a cue conflict—forward displacement of the visual target from the proprioceptive target—there is the potential for a confound related to visual recalibration. During cue conflict blocks, the white square shifted forward on both VP and V targets, the latter of which were used to assess visual recalibration. Ideally, participants should place their indicator finger at the perceived location of the white square on V trials, whether shifted or not. If they undershoot by the end of the cue conflict block relative to the beginning of the block, we infer that they perceive the visual target to be closer than it actually is (visual recalibration). The potential confound is that participants might have a general tendency to avoid moving their indicator hand farther away, i.e. effort minimization. This could result in undershooting the visual target as it gets farther away.

To control for this possibility, participants performed a behavioral experiment identical to the perceptual alignment task in Expt. 2, except that the “cue conflict” block consisted only of V trials. The white box shifted forward at the same rate as the main experiments, reaching 70 mm forward displacement after 84 trials. Participants were trained on all three target types as in the main experiments, but there were no P or VP trials during the V target shift, so there was no cue conflict. Thus, we did not expect visual recalibration (undershoot) unless participants were minimizing movement distance of the indicator hand. Visual recalibration was calculated the same as the main experiments (equation 2).

### Statistical Analysis

We tested whether visual recalibration differed significantly from zero using a one-sample t-test. For context, we applied the same test to visual recalibration in the cue conflict and veridical sessions of Expt. 1 (expected to be greater than zero and similar to zero, respectively) and in Expt. 2 sham group (expected to be greater than zero). In each case, the test was performed two-tailed with alpha of 0.05. Cohen’s d was computed for effect size.

## RESULTS

### Experiment 1

#### Behavior

Each participant completed both a cue conflict session and a veridical session, on different days in random order (Fig. 3A). In the cue conflict session, the visual cue was gradually shifted forward of the proprioceptive cue over the course of 84 trials, to a max of 70 mm. On average, participants recalibrated vision 34.7 ± 6.9 mm (mean ± 95% CI) and proprioception 11.1 ± 5.3 mm (Fig. 3Di). Some participants recalibrated proprioception more than vision (Fig. 3Bi) while others recalibrated vision more than proprioception (Fig. 3Ci). As shown previously, visual and proprioceptive recalibration were inversely associated (r = -0.52, p = 0.013) (Fig. 3Diii)^9^.

In the veridical session, there was no cross-sensory mismatch, so the amount of visual and proprioceptive recalibration is expected to be roughly zero on average. In line with previous work, participants recalibrated vision -1.2 ± 5.0 mm and proprioception -2.2 ± 4.9 mm (Fig. 3Dii)^9,10^.

### Neurophysiology

We found no evidence that baseline neurophysiological measures differed across sessions (Table 1), or that SAI changed from pre-alignment task to post-alignment task timepoint in either session (ANOVA Session x Timepoint: F_1,21_ = 1.08, p = 0.31; Session: F_1,21_ = 3.70, p = 0.068; Timepoint: F_1,21_ = 2.29, p = 0.15) (Fig. 3E). We also found no change over time in unconditioned motor evoked potential (MEP) amplitude (Session x Timepoint: F_1,21_ = 0.31, p = 0.58; Session: F_1,21_ = 0.08, p = 0.78; Timepoint: F_1,21_ = 2.72, p = 0.11), indicating that any associations between SAI and recalibration behavior were not confounded by differences in motor cortex excitability.

**Table 1.** Experiment 1 summary of neurophysiological measures recorded at baseline in each session and subjective ratings recorded at the end of each session. None differed across sessions (all p > 0.2).

| | **Cue Conflict Session**<br>(mean ± 95% CI) | **Veridical Session**<br>(mean ± 95% CI) |
| --- | --- | --- |
| RMT (% MSO) | $38.5 \pm 2.5\%$ | $38.5 \pm 3.0\%$ |
| SI_1mV (MEP in mV) | $1.0 \pm 0.2$ mV | $1.0 \pm 0.1$ mV |
| SAI (% of unconditioned MEP) | $58.5 \pm 10.4\%$ | $50.0 \pm 11.3\%$ |
| Attention rating | $7.5 \pm 0.4$ | $8.0 \pm 0.3$ |
| Fatigue rating | $4.4 \pm 0.7$ | $3.4 \pm 0.8$ |
| Sleep rating | $7.7 \pm 0.5$ | $7.4 \pm 0.5$ |
| Pain rating | $2.6 \pm 0.8$ | $2.8 \pm 0.7$ |

RMT: Resting motor threshold. MSO: Maximum stimulator output. SI\_1mV: motor evoked potential (MEP) achieved at a stimulus intensity intended to elicit a 1mV MEP. SAI: Short latency afferent inhibition. All ratings were on a scale from 1-10, reported verbally, with 10 being the best attention and sleep and the worst fatigue and pain.

Two example participants with different magnitudes of proprioceptive recalibration in the cue conflict session are shown in Figure 3. The individual with relatively larger proprioceptive recalibration had a decrease in SAI post-cue conflict (Fig. 3Bii), whereas the individual with smaller proprioceptive recalibration had an increase in SAI post-cue conflict (Fig. 3Cii). Consistent with this pattern, multilevel regression model results suggest that change in SAI from before to after the alignment task (post divided by pre) showed modality-specific recalibration associations with the cue conflict, but not the veridical session (Table 2). These associations are illustrated with predictor residual plots (Fig. 3F), which allows us to show the relationship of one predictor variable with the dependent variable, after statistically controlling for the effect of the other predictor^50^. For either modality, recalibration in the positive direction is beneficial in the cue conflict session (helps compensate for the visuo-proprioceptive mismatch).

**Table 2.**
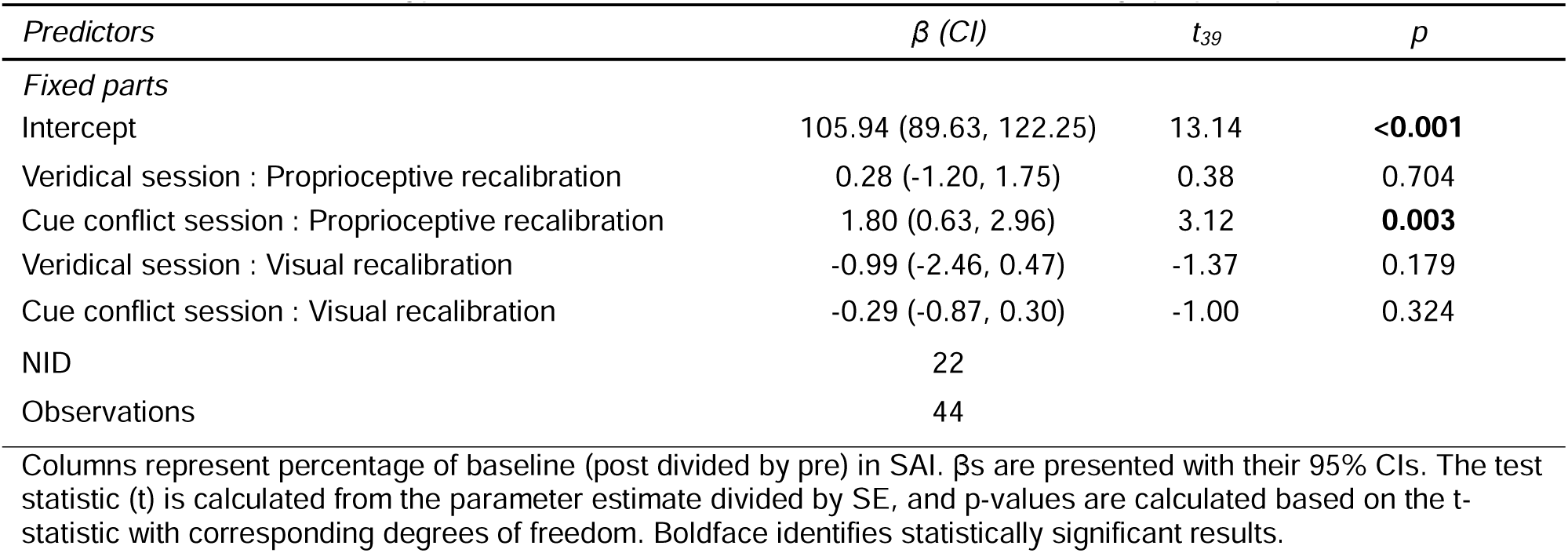
Experiment 1 multilevel regression results for short latency afferent inhibition (SAI), comprising four interaction terms for session type (Veridical or Cue conflict) and recalibration modality (proprioceptive or visual).

After controlling for the effect of visual recalibration, greater positive proprioceptive recalibration was associated with greater decrease in SAI (less inhibition post-alignment task) in the cue conflict session (β = 1.80, t_39_ = 3.12, p = 0.003), but not the veridical session (β = 0.28, t_39_ = 0.38, p = 0.704) (Fig. 3Fi). After controlling for the effect of proprioceptive recalibration, change in SAI was not significantly associated with visual recalibration (cue conflict session: β = -0.29, t_39_ = -1.00, p = 0.324; veridical session: β = -0.99, t_39_ = -1.37, p = 0.179; Fig. 3Fii). This supports the prediction in the upper right (orange) cell in Fig. 1C.

In the veridical session, SAI at baseline was negatively correlated with proprioceptive variance (r = -0.45, p = 0.034). Baseline SAI was not significantly correlated with visual variance (r = - 0.11, p = 0.64) or with participants’ weighting of vision vs. proprioception on bimodal trials (r = 0.18, p = 0.42). In the cue conflict session, SAI at baseline was not significantly correlated with proprioceptive recalibration (r = -0.19, p = 0.40).

### Experiment 2

#### Veridical blocks

Three groups of 27 participants performed a block of V, P, and veridical VP trials before and after cTBS was delivered over their target hand representation in SI, M1, or sham (Fig. 4A). We computed 2D variance of participants’ estimation endpoints on unimodal V and P trials in each veridical block to assess whether visual or proprioceptive variance changed after cTBS (Fig. 4B). Proprioceptive variance was similar across groups before cTBS (M1: 56.1 ± 16.6; SI: 53.8 ± 12.7; Sham: 60.5 ± 14.4 mm^2^) (mean ± 95% CI). Proprioceptive variance changed differently across groups post cTBS relative to pre, with the difference between pre and post showing an effect of group (ANOVA: F_2,78_ = 4.31, p = 0.017, η^2^ = 0.10). This was driven by an increase in proprioceptive variance post SI cTBS relative to sham (p=0.013) (Fig. 4Ci). Visual variance was similar across groups before cTBS (M1: 72.2 ± 17.0; SI: 91.6 ± 30.7; Sham: 68.5 ± 28.9 mm^2^) (mean ± 95% CI). cTBS had no impact on visual target estimation variance regardless of group (χ^2^ = 2.08, p = 0.35) (Figure 4Cii). This was also the case when the analysis was performed without the two outliers(p > 0.2).

We also computed an estimate of how much people were relying on vision vs. proprioception when both were available (weighting), taking advantage of subjects’ naturally different spatial biases when estimating V, P, and VP targets^51,52^, even with no cue conflict^31^. As computed, this parameter ranges from 0 (only proprioception) to 1 (only vision), with 0.5 indicating equal reliance on vision and proprioception. Weighting was similar across groups before cTBS, with values in line with past work^32,35^ (M1: 0.39 ± 0.07; SI: 0.46 ± 0.09; Sham: 0.41 ± 0.09) (mean ± 95% CI). cTBS had no impact on weighting regardless of group (F_2,76_ = 0.308, p = 0.74, η^2^ = 0.008).

### Cue conflict block

On average, visual recalibration was 36.8 ± 7.2 mm for M1, 38.8 ± 7.7 mm for SI, and 32.7 ± 8.4 mm for Sham (mean ± 95% CI). Proprioceptive recalibration was 6.5 ± 6.0 cm for M1, 18.1 ± 7.0 cm for SI, and 9.6 ± 4.5 mm for Sham (Fig. 5A-B). Proprioceptive and visual recalibration were inversely correlated for all groups; individuals with larger proprioceptive recalibration had smaller visual recalibration and vice versa (M1: r = -0.57, p = 0.002, SI: r = -0.65, p < 0.001, Sham: r = -0.40, p = 0.04) (Fig. 5C).

**Figure. 5.**
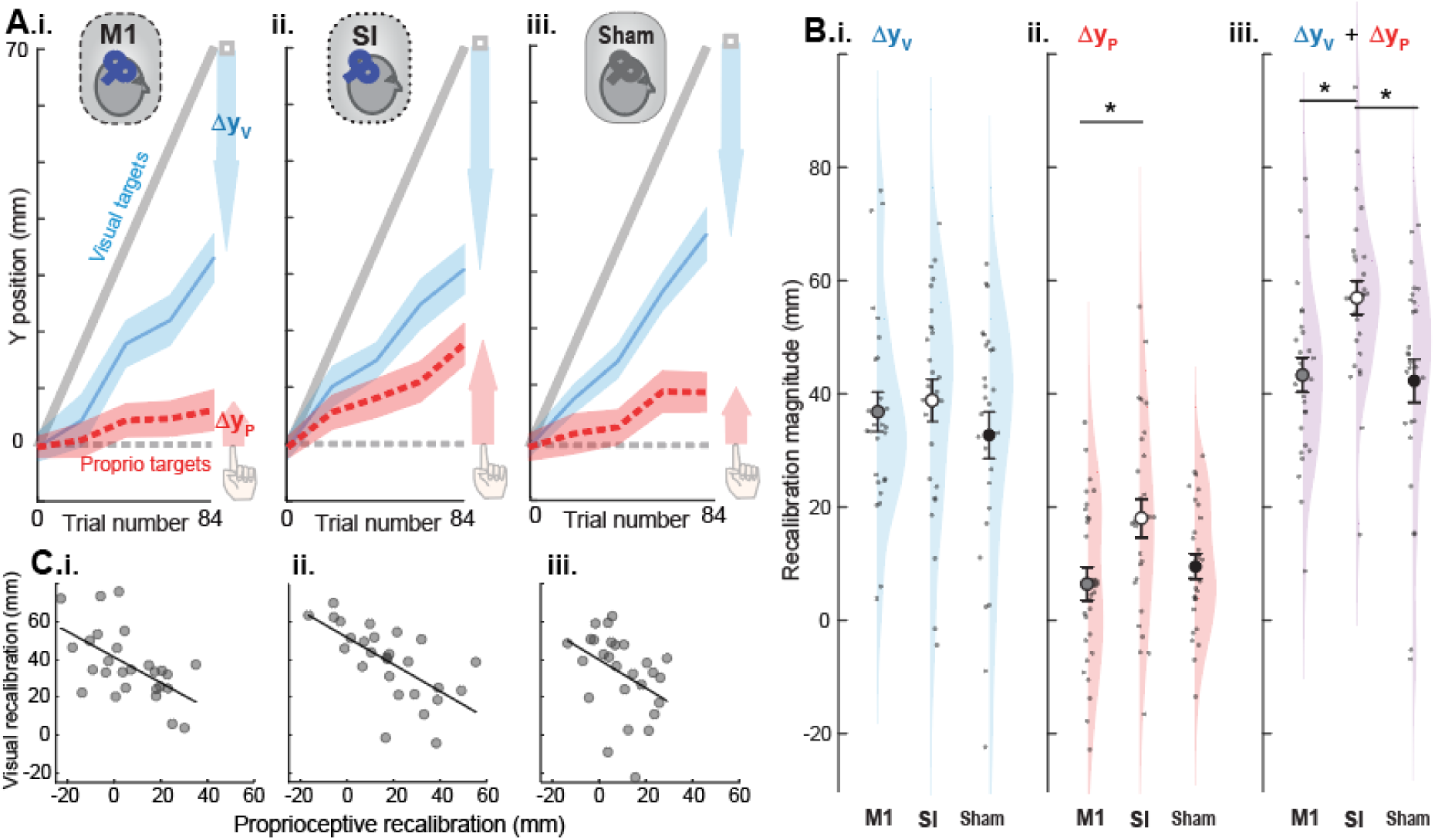
Experiment 2 group behavior during the cue conflict block. **A.i-iii.** Mean performance on P and V targets in the cue conflict block after M1, SI, and Sham cTBS, respectively (averaged every 4 trials for clarity). Undershooting the visual target on V trials represents visual recalibration (blue arrow) and overshooting the proprioceptive target on P trials represents proprioceptive recalibration (red arrow). SI participants recalibrated proprioception noticeably more than the other groups. Shading represents SEM. **B**. Comparing recalibration across groups. **i.** Visual recalibration did not differ significantly across groups **ii.** Proprioceptive recalibration was larger in the SI group compared to the M1 group. **iii.** Total recalibration (sum of visual plus proprioceptive recalibration) for SI was greater than M1 or Sham groups. **C. i-iii**. Across all groups, proprioceptive recalibration was negatively associated with visual recalibration.

While visual recalibration was similar across groups (F_2_,_78_=0.68, p=0.51, η_p_^2^= 0.017) (Figure 5Bi), the magnitude of proprioceptive recalibration (Figure 5Bii) and total recalibration (Figure 5Biii) differed across groups (F_2_,_78_=4.35, p=0.016, η_p_^2^ = 0.100; F_2_,_78_=6.18, p=0.003, η_p_^2^ = 0.137). Proprioceptive recalibration was larger following SI cTBS relative to M1 cTBS (p=0.015) but not Sham (p=0.099). In contrast, proprioceptive recalibration was similar between M1 and Sham (p=0.73). Total recalibration (sum of visual plus proprioceptive recalibration, reflecting the total magnitude participants compensated for the perturbation) was larger following SI cTBS compared to M1 or Sham (p=0.012, p=0.006, respectively). However, there was no difference in total recalibration between M1 and Sham (p=0.97). Together, these results suggest that SI cTBS had an effect on how participants compensated for the cue conflict, and that this effect was driven by proprioceptive recalibration. This supports the prediction in the right column of Fig. 1C.

### Subjective ratings and tactile sensitivity

Subjective ratings of pain, sleep quality, attention, and fatigue were similar across groups, as was baseline neurophysiology (Table 3).

**Table 3.**
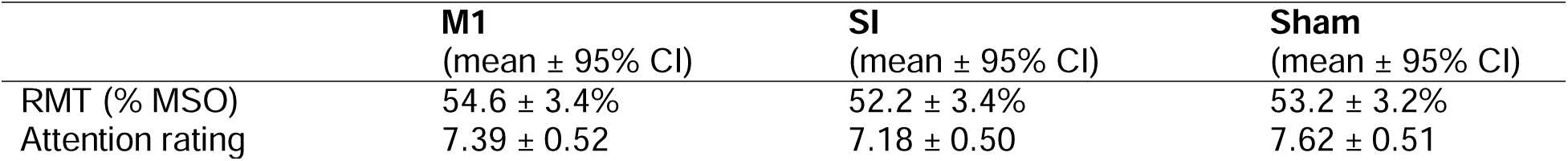
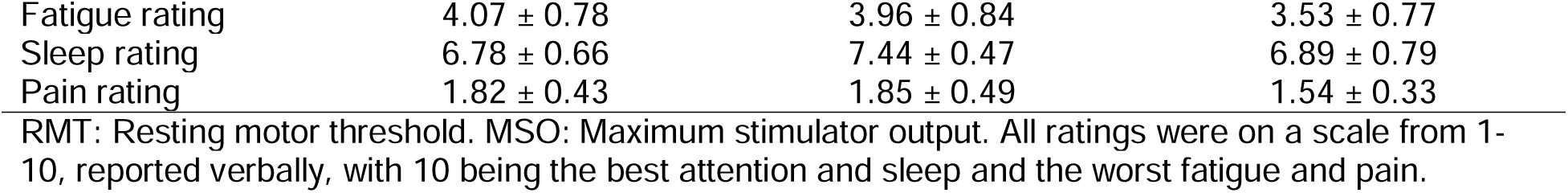
Experiment 2 summary of neurophysiology and subjective ratings across groups.

An adapted grating orientation test (GOT) was used as a rough estimate of tactile sensitivity in the target index finger before and after cTBS. After cTBS, the M1 and SI groups showed a change in GOT threshold of -0.04 ± 0.26 and 0.12 ± 0.27, respectively (mean ± 95% CI). The sham group change in GOT threshold was 0.004 ± 0.24. Change in GOT after cTBS did not differ significantly across groups (χ^2^ = 0.33, p = 0.85). Bayes Factor was calculated as 0.148, denoting substantial evidence for equivalence among the groups.

### Control Experiment

We hypothesized that in the absence of bimodal VP trials to create a cue conflict there would be no visual recalibration, and therefore subjects would closely follow the visual cue without undershooting it. Participants did not have any apparent difficulty pointing consistently at the V target throughout the gradual 70 mm displacement (Fig. 6A). The computed average change in V target undershooting was -2.32 ± 5.52 mm (mean ± SE), which was not significantly different from zero (t_17_ = -0.42, p = 0.68, Cohen’s d = -0.099). Together, these results are consistent with no visual recalibration, or minimization of effort, on average, despite the forward 70 mm shift of the visual target.

**Fig. 6.**
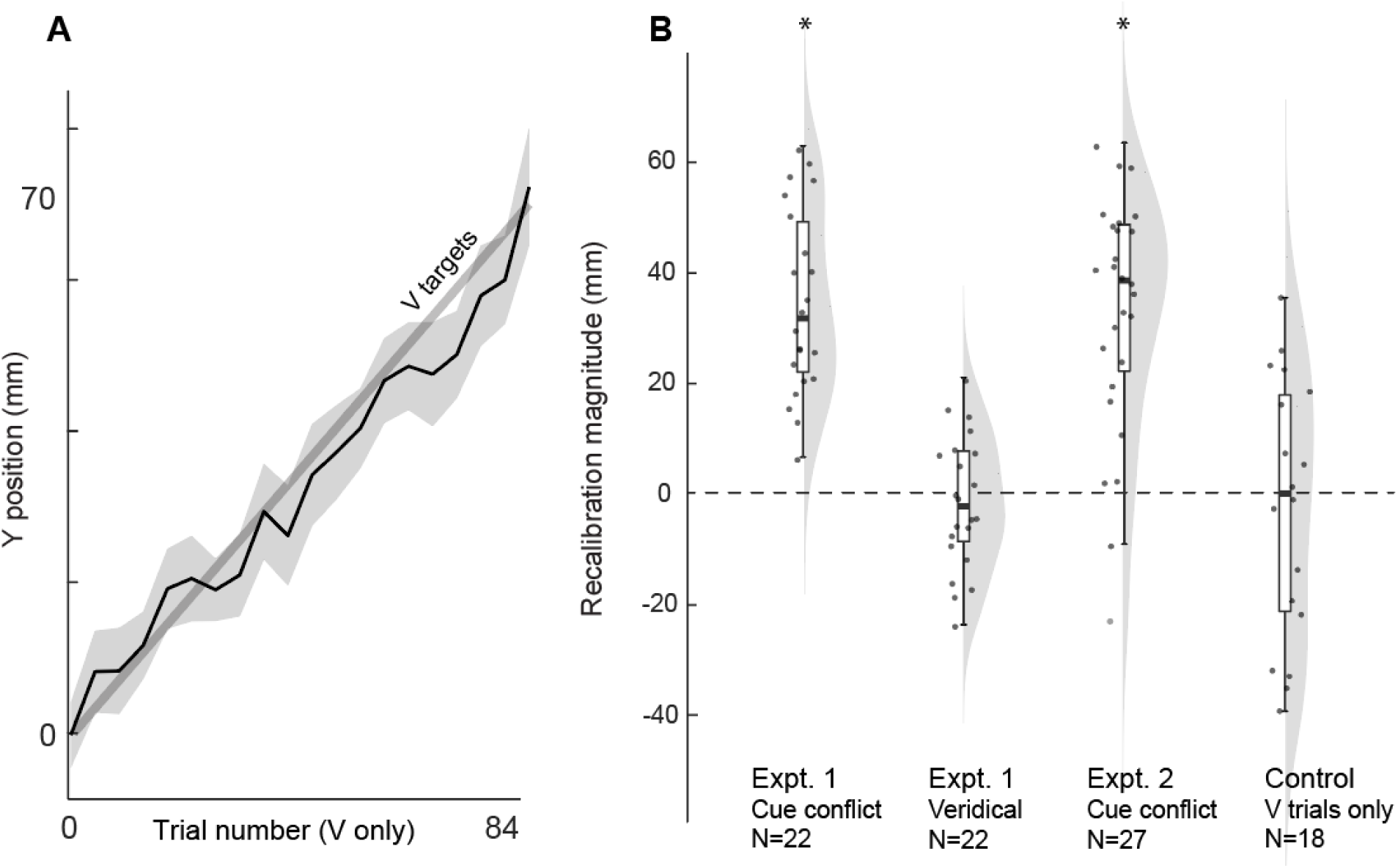
Control experiment results. **A.** In the absence of P or VP trials, participants successfully pointed (black line) at the visual target (thick grey line) throughout the 70 mm displacement (averaged every 4 trials for clarity). Shading represents SEM. **B.** Visual recalibration computed for the Expt. 1 cue conflict and veridical sessions (expected to be greater than and similar to zero, respectively), the Expt. 2 sham group cue conflict block (expected to be greater than zero), and the control experiment (expected to be similar to zero). Single subjects (dots), box plots (white bars), and violin plots (grey shading) are shown to illustrate each distribution. *p<0.05.

We also compared visual recalibration in the control experiment with how this parameter looked when there actually was a cue conflict in the two main experiments (Fig. 6B). As expected, visual recalibration was significantly greater than zero in the cue conflict session of Expt. 1 (t_21_ = 9.80, p < 0.001, Cohen’s d = 2.09) and in the cue conflict block of Expt. 2 (t_26_ = 7.96, p < 0.001, Cohen’s d = 1.53). Thus, in the presence of a 70 mm forward shift of the visual cue relative to the proprioceptive cue in bimodal trials, visual recalibration is large and easily detectable.

Finally, we examined visual recalibration in the veridical session of Expt. 1 (Fig. 6B). As expected, in the absence of any cue conflict or shift of the visual target, there was no significant visual recalibration (t_21_ = -0.49, p = 0.63, Cohen’s d = -0.104). Descriptively, we notice that the distribution of visual recalibration is less spread out for the veridical session of Expt. 1 than for any of the comparisons, including the control experiment. This could suggest that shifting the visual target adds noise, even if the response is unbiased across subjects.

In sum, neither bimodal targets (veridical session) nor a forward-shifted visual cue (control experiment) was sufficient on its own to induce nonzero visual recalibration; this only occurred in response to a forward-shifted visual cue combined with bimodal targets, as in the cue conflict conditions.

## DISCUSSION

We began with two questions: (1) Does SI convey hand representation updates to the motor system? (Fig. 1C rows) and (2) Is this involvement multisensory or proprioception-specific? (Fig. 1C columns). Experiment 1 showed that SAI, a measure of somatosensory-motor integration, changed with proprioceptive, but not visual, recalibration, and only in the presence of a cue conflict. Experiment 2 found that modulating SI, but not M1, with cTBS increased proprioceptive variance and recalibration, without the same effect on visual responses. These findings suggest that in the presence of a visuo-proprioceptive cue conflict, SI conveys proprioceptive hand representation updates to M1 (Fig. 1C orange cell).

### Plasticity in SI projections to M1 reflects proprioceptive recalibration

In Experiment 1, changes in SAI were associated with proprioceptive recalibration in the cue conflict session but not the veridical session. We previously found that individuals who recalibrated proprioception more had larger decreases in M1 excitability, while individuals who recalibrated vision more had larger increases in M1 excitability^9,53^. Since the magnitude of SAI is proportional to the magnitude of somatosensory afference to SI^54^, decreased SAI likely reflects decreased SI excitability, which may have driven the previously observed decreases in M1 excitability in individuals who recalibrated proprioception to a greater extent^9,10^. In contrast, the lack of association between SAI changes and visual recalibration suggests that changes in M1 excitability that we previously associated with visual recalibration are mediated by areas outside of SI, such as multisensory posterior parietal or premotor areas^22,24,55–57^.

Decreased somatosensory projections to motor cortex may reflect somatosensory gating to help resolve a multisensory conflict. Decreased SAI has also been reported after the rubber hand illusion (RHI), another multisensory cue conflict paradigm^19^. Contrary to the RHI, we found no average difference in SAI after the visuo-proprioceptive cue mismatch relative to veridical session. This may be explained by differences in these two paradigms. While our visuo-proprioceptive task and RHI both involve a multisensory conflict, the RHI involves synchronous stroking of a seen fake arm and felt real arm, which depends not only on spatial visuo-proprioceptive cues but also temporal visuo-tactile cues. In addition, the subject knows consciously the whole time that the fake arm is not their real arm^58,59^. In contrast, participants are mostly unaware of the visuo-proprioceptive conflict in the present study^36^.

Besides using SAI to explore plasticity in somatosensory projections to M1 in response to a visuo-proprioceptive cue conflict, baseline SAI may have functional relevance that predicts how individuals respond during the visuo-proprioceptive task. We found that baseline SAI was associated with proprioceptive variance, but not visual variance, weighting, or recalibration, suggesting that baseline SAI may reflect a more low-level proprioceptive process rather than the computations involved in multisensory integration during a cue mismatch. The lack of association between baseline SAI and recalibration is similar to that observed with the RHI^19^.

Instead, baseline long latency afferent inhibition was associated with the strength of the illusion, suggesting that processing in other areas like premotor and parietal cortices^26,56,60^ may be important in understanding mechanisms of multisensory integration.

### SI activity is important for proprioceptive variance and recalibration

If previously reported modality-specific associations between recalibration and M1 excitability^9,10^ were driven by somatosensory projections to M1, then one would predict dissociable effects of M1 and SI neuromodulation on visuo-proprioceptive recalibration. To test this, in Experiment 2 we delivered cTBS over M1 or SI or sham, and quantified changes in veridical task performance and on recalibration in response to a cue conflict. SI, but not M1, cTBS affected proprioceptive variance, supporting the role of SI in proprioceptive processing for guiding action.

SI cTBS increased total recalibration, a difference apparently driven by proprioceptive rather than visual recalibration. However, SI cTBS did not disrupt the normal inverse relationship between proprioceptive and visual recalibration^2,12–14^. This is consistent with SI’s role in proprioceptive recalibration being somewhat independent of other multisensory computations that mediate this relationship, such as parietal^14,24,55^ or prefrontal cortices^61,62^, or the cerebellum^63^. In other words, disrupting SI did not evidently disrupt the coordination of the visual vs. proprioception relationship in responding to a cue conflict. Even theoretically “unisensory” areas such as SI are involved in cross-sensory interactions^64^, and our results do not preclude SI being involved in, or having access to, integrated multisensory information about the hand.

However, the cue conflict-related signals sent by SI to M1 appear limited to proprioception.

The increase in proprioceptive variance and recalibration associated with SI cTBS raises questions about potential changes in tactile function, as participants relied on feeling smooth or rough markers to place their target finger. However, SI cTBS did not affect the grating orientation test (GOT), indicating no confounding impact on tactile function. While prior studies reported SI cTBS impairing tactile perception, these used small samples and a variety of methods and timing. Given the depth and focality limits of TMS^45^, it likely stimulated both tactile and proprioceptive regions within SI^46^. Importantly, the effects of SI and M1 cTBS were dissociable on a multisensory perceptual task that guided action, the main focus of this study.

### Motor system considerations

Proprioception plays an integral role in motor control, and M1, the cortical region most associated with movement execution, is closely interconnected with SI. The neural basis of these reciprocal connections has been supported behaviorally; a visuo-proprioceptive cue conflict in the absence of motor adaptation not only affects perception, but also subsequent reaching movements, suggesting a common sensorimotor map for perception and action in this paradigm^11^. In other words, any changes in perceived hand position need to be considered when the brain plans hand movements. This presumably reflects a change in SI, M1, or both. It is therefore interesting to note that SAI changed in association with proprioceptive recalibration, and SI but not M1 cTBS increased proprioceptive recalibration; this is consistent with the idea that motor effects of proprioceptive recalibration may be mediated by the SI-to-M1 pathway, although not directly controlled by M1 itself.

Given SI’s involvement in motor planning^65^ and skill learning^38^, we must ask if our findings could reflect motor processing rather than proprioceptive recalibration. Proprioceptive recalibration often occurs in visuomotor adaptation, where sensory prediction errors and visuo-proprioceptive cue conflict drive adjustments^1^. However, visuomotor adaptation is unlikely here, as participants received no online or endpoint movement feedback. While the offset visual cue after placing the target finger on a tactile marker might resemble a sensory prediction error, this is unlikely since (1) the tactile marker itself was the explicit movement goal, and (2) our prior studies indicate subjects have low certainty about the visuo-proprioceptive offset in this paradigm^36,66^.

Our results align with motor learning research that suggests SI and M1 play distinct roles in motor adaptation and retention^67,68^. Suppressing SI, but not M1, has been shown to impair adaptation without affecting movement kinematics, indicating SI’s specific role in learning mechanisms^67^. SI is thought to contribute to the encoding and retention of learned movements^67–69^, motor learning by observation^70,71^, and changes in proprioception accompanying motor learning^17^. If what is learned in such tasks is partly an updating of the hand representation, our findings could clarify SI’s contribution to learning. SI cTBS effects on proprioceptive variance and recalibration suggest that SI’s role in motor adaptation may stem from changes in limb representation influencing motor learning.

### Visual recalibration does not reflect effort minimization

Previous studies have demonstrated both visual and proprioceptive recalibration in response to a cue conflict^72,73^, though some have suggested visual recalibration is minimal^74,75^. Visual recalibration likely depends on the weighting of visual cues, which varies by task. In the present cue conflict paradigm, participants rely more on proprioception than vision^35^, consistent with evidence that recalibration is inversely related to cue weighting. However, we wanted to test the possibility that what we refer to as visual recalibration (increasing undershoot of the visual target) is not an artifact of introducing the cue conflict by a forward displacement of the visual cue. In other words, participants might have a general tendency to avoid moving their indicator hand farther away as the visual cue shifts, i.e. effort minimization rather than a change in perception.

The control experiment tested this with a block of visual cues gradually shifting forward, with no bimodal cues to induce a cue conflict. Participants shifted their estimates along with the shifting cue, with no increase in undershooting. Visual cue undershooting was specific to the cue conflict condition, which is inconsistent with the idea of effort minimization.

## Conclusions

Representation of the hand is critical for accurate control of movement, and our results suggest that SI plays a key role in updating the motor system when this representation is changed by a cue conflict. While SI’s role was specific to the presence of a visuo-proprioceptive cue conflict, SI activity conveyed to M1 reflects proprioceptive and not multisensory recalibration. These findings add to our understanding of the interaction of multisensory body representation with control of movement.

## Author Contributions

JLM: Methodology, Formal Analysis, Investigation, Writing – Original Draft, Writing – Review & Editing, Visualization. RB, MW, & TLM: Methodology, Formal Analysis, Investigation. CRS & AH: Formal Analysis, Investigation. HJB: Conceptualization, Methodology, Software, Writing – Original Draft, Writing – Review & Editing, Visualization, Supervision, Funding Acquisition.

## Competing Interest Statement

The authors have no competing interests to disclose.

## ACKNOWLEDGMENTS

The authors wish to acknowledge Indiana University Biostatistics Consulting Center; specifically, Lilian Golzarri Arroyo for assistance with the hierarchical linear regression analysis in Expt. 1, and Stephanie Dickinson for randomizing group assignment in Expt. 2.

## FUNDING

National Institute of Neurological Disorders & Stroke (NINDS) grant R01 NS112367 to HJB.

## Data Availability

The data and analysis code used to produce this manuscript are publicly available at https://osf.io/w5pfk/.

## Notes

https://osf.io/w5pfk/

### Competing Interest Statement

The authors have declared no competing interest.

### Summary of Updates

The methods section has been expanded to provide more detail on the behavioral task and the reasoning behind specific methodological choices. The introduction has been revised, and a new conceptual figure added, to better explain how specific results support the conclusions reached. A control experiment has been added to resolve the question of whether participants undershoot the visual cue because they are minimizing their physical effort (a confound) or because they are recalibrating their visual estimate of its location. The control experiment clearly contradicts the idea of effort minimization. Other changes to the presentation of the data and methods have been made to improve clarity.

